# Assessing the Rate-Dependence of the First Phase of Glucose-Stimulated Insulin Secretion: Dynamic Perifusion Studies with Isolated Human Pancreatic Islets

**DOI:** 10.1101/2025.07.01.662581

**Authors:** Peter Buchwald, Sung-Ting Chuang, Brandon Watts, Oscar Alcazar

## Abstract

Insulin released in response to a stepwise increase in glucose (square wave) is biphasic with a transient first-phase peak of 5-10 min and a more sustained second phase. While it is usually assumed that the first phase is rate- and the second is concentration-dependent, there are no detailed investigations into the rate-sensitivity of the first phase. We performed dynamic perifusion studies with isolated human pancreatic islets in a system that allowed fully customizable glucose ramps and established the corresponding insulin secretion time-profiles with high (once-a-minute) time-resolution. We considered the first-phase release to be the excess amount of insulin in addition to what would be expected from the glucose concentration-dependent second-phase release and examined its dependence on the glucose gradient (rate of increase). The average first-phase insulin release rate calculated this way increased with the gradient and could be fitted well with a Hill-type sigmoid function having a half-maximal value around 1.25 mM/min (*n*_Hill_=1.8, *r*^2^=0.96). This agrees well with our previously introduced glucose-insulin control system built on a general framework of a sigmoid proportional–integral–derivative (σPID) controller, a generalized PID controller more suitable for biological systems than classic linear ones as biological responses are always limited between zero and a possible maximum. Experimental results obtained here were used to slightly readjust the parameters of this local glucose concentration-based computational model resulting in predictions in good agreement with measured first- and second-phase insulin secretions (*r*^2^>0.90). Thus, glucose-stimulated insulin secretion (GSIS) of perifused human islets can be described well as the sum of a rate-sensitive first phase, which is a sigmoid function of the glucose gradient with half-maximal value around 1.25 mM/min, and a concentration-sensitive second phase, which is a sigmoid function of the glucose concentration with half-maximal value around 8 mM.

## Introduction

Glucose levels are maintained in their normal range almost exclusively by the insulin released from the beta cells located in pancreatic islets. If blood glucose in humans rises above ∼11 mM, it passes the renal filtration threshold resulting in too much glucose being eliminated into the urine, which drags water osmotically and leads to secretion of large volumes of sweet urine, the classic symptom of diabetes [1]. Therefore, a highly sensitive and fine-tuned glucose-insulin control system is needed to maintain the blood glucose in its relatively narrow normal range around the glycemic set-point – about 3 to 9 millimolar (mM) in humans (corresponding to 54 to 162 mg/dL) with normoglycemia at ∼5 mM (90 mg/dL) [1]. Pancreatic islets seem to be capable of achieving this independently by acting as glucose sensors and regulating their insulin secretion accordingly [1]; for example, they can maintain their own glycemic set point when transplanted into different species [2].

Accurate quantitative characterization of glucose-stimulated insulin secretion (GSIS) is of considerable interest not only for describing the normal physiological mechanism, but also for assessing β-cell function, as abnormalities are critical in both type 1 and type 2 diabetes (T1D, T2D), and for the development and improvement of artificial/bioartificial pancreas systems including automated closed-loop insulin delivery devices [3] as well as immune-isolated, encapsulated islets [4-7]. It has been known since the late 1960s that insulin released in response to an abrupt stepwise increase in glucose (square wave) is biphasic with a transient 5–10 min long first-phase peak followed by a more sustained second phase [8-13], an observation confirmed and characterized multiple times in different species with isolated islets (Figure 1), isolated pancreases, and *in vivo* [8, 14-18]. In fact, more generally, there are four or possibly even five modes / phases of insulin secretion [19]: (1) basal insulin secretion corresponding to release in the postabsorptive state; (2) cephalic insulin secretion, which is evoked by the sight, smell, and taste of food even before any nutrient is absorbed and is mediated by pancreatic innervation; (3) first-phase insulin secretion – the initial burst released in the first 5–10 min after exposure to a rapid increase in glucose (or other secretagogues); (4) second-phase insulin secretion, which follows the acute response, is related to the degree and duration of the stimulus, and is flat or slowly rising; and (5) possibly a third phase that has been described but only *in vitro*.

**Figure 1.**
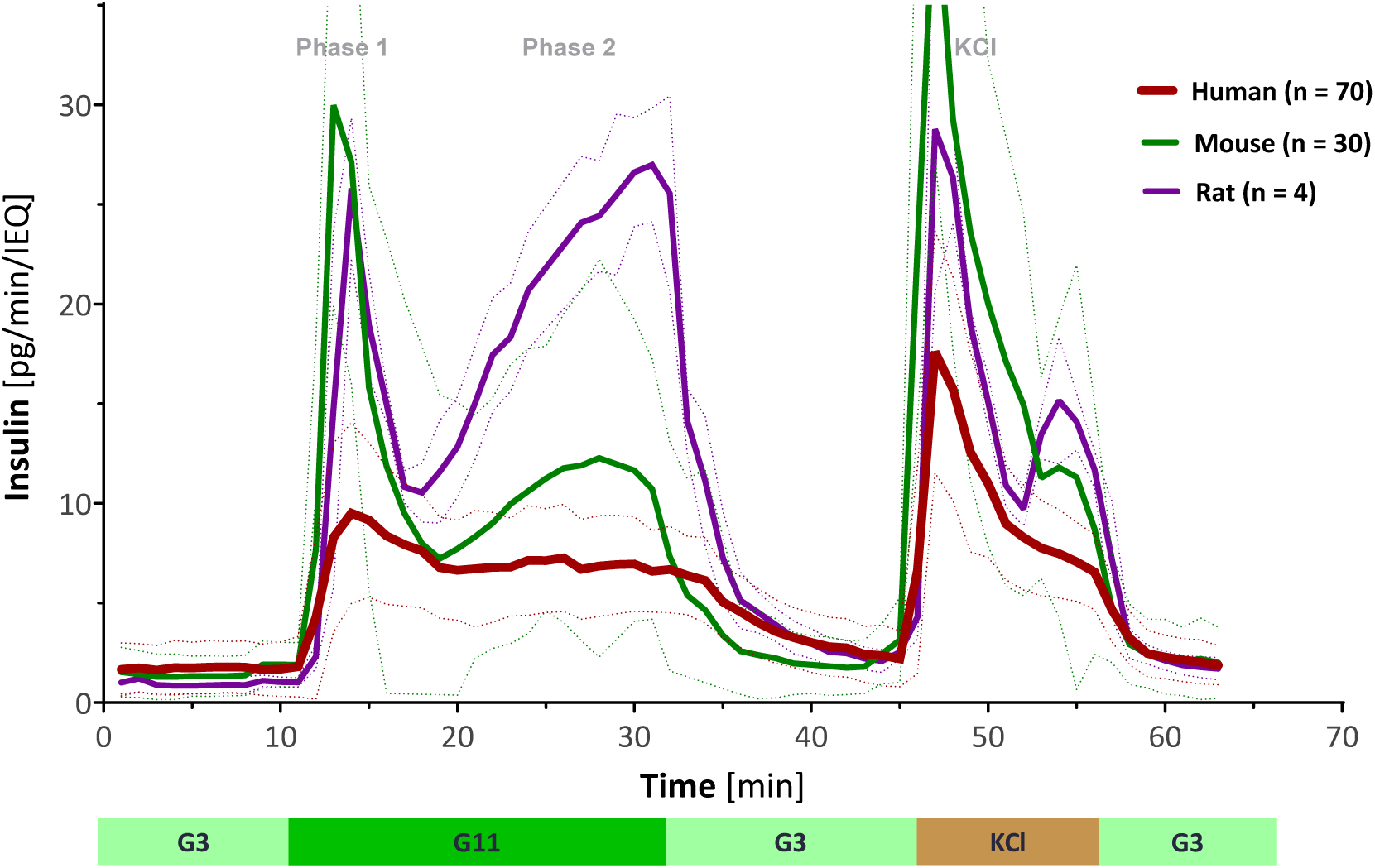
Insulin secretion rates in perifused human, mouse, and rat islets (dynamic GSIS). Average of multiple experimental data collected for untreated control islets perifused using an automated multichannel perifusion apparatus and a standard protocol of low (3 mM; G3, 8 min), high (11 mM; G11, 20 min), and low (3 mM; G3, 15 min) glucose stimulation (followed by KCl and G3, each for 10 min) as shown (output samples collected every minute; 0.1 mL/min flow rate, ∼100 IEQ per channel). Data are average ± SD from multiple isolations with number of samples (*n*) for each species indicated in the graph (SDs shown as thinner dashed lines of the same color). While rat data were obtained on a much more limited sample size (*n* = 4) than human or mouse data (*n* = 70 and 30, respectively), they were included here as their overall shape agrees very well with that obtained in other larger studies (e.g., [55]).

Regarding the biphasic response, it is typically assumed that the more sustained second-phase release is related to the glucose level (concentration, *c*_g_) whereas the burst-like first-phase release is triggered by quick increases in glucose, thus, it is likely related to the rate of change in glucose (i.e., its time gradient ∂*c*_g_/∂*t*). The latter can provide a sort of differential control, which is usually included in control systems to provide a mechanism for quick responses to changes [1]. The presence of a biphasic pattern is considered an important indication of adequate β-cell function [13, 19-21], and loss of the first-phase response has been shown to be a possible early sign of progress toward both T1D [22-26] and T2D [13] (and references therein). While the concentration-dependency of insulin secretion and in particular its second phase have been characterized in detail, including relatively recently by our group [18], there seem to be no detailed investigations, especially not with human islets, into the rate sensitivity of the first-phase response. Thus, it has not been established how fast the transition has to be to trigger a noticeable first phase or how does the amount of insulin released in this first phase depend on the abruptness and duration of this increase (e.g., the slope and duration of the glucose ramp). Toward this aim, here, we performed dynamic perifusion studies with isolated human pancreatic islets in a system that allowed custom adjustable transition times for the increase in incoming glucose level and established the corresponding insulin secretion time-profiles with high (i.e., once a minute) time-resolution.

Perifusion studies of isolated islets, which have been around since the 1970s [27-29] and have improved considerably due to technical and analytical advances, allow the detailed mapping of the time profile of insulin release under fully controllable influxes of glucose, oxygen, and other secretagogues of interest. Due to advances in microfluidics, even the monitoring of insulin secretion of single islets is now possible [30-32]. Perifusion studies represent the most complex *in vitro* assay to assess the quality and function of isolated islets or other insulin secreting cells and provide considerably more information than the extensively used static GSIS and corresponding stimulation indices (SIs). As islets are mini-organs, isolated islets reproduce the function of those in their native physiological environment quite closely; thus, detailed investigations can be performed to obtain data that otherwise could not be obtained *in vivo*. There are potentially some metabolic limitations with isolated islets due to loss of vasculature, hypoxia being the most critical one. However, due to the flowing media, these limitations are much less severe in dynamic perifusion than in static incubation assays. This could very well be a main reason why stimulated insulin secretion in perifusion assays tend to not correlate well with static culture SIs, as it has been shown, for example, in a relatively large analysis of human islets isolated at multiple US centers [33]. Static incubation systems are also prone to accumulation of excretion metabolites, some of which might further hamper GSIS responses in such systems (e.g., lactic acid accumulation may alter the medium pH).

Here, we performed dynamic perifusion studies using various constant glucose ramps with transition times increasing up to 20 min. We considered the first-phase release to be the excess amount of insulin in addition to what would be expected from the glucose concentration-dependent second-phase release and plotted it as a function of the rate of glucose increase to establish its rate-dependence. Finally, we fitted both the first-phase response as calibrated here and the second-phase response whose concentration-dependence we calibrated earlier [18] with our previously introduced computational model of glucose-insulin control system built on a general framework of a sigmoid proportional– integral–derivative (σPID) controller [34, 35] and tested its ability to predict insulin responses generated by the various incoming glucose influx conditions.

## Methods

### Human islets

Human pancreatic islet samples were procured from the Integrated Islet Distribution Program (IIDP) at City of Hope (Duarte, CA, USA). All islet samples used here were from non-diabetic donors; characteristics of the human islet donors for the present study are summarized using standard checklists recommended for reporting human islet preparations used in research in Supplementary Material Table S1.

### Islet perifusion

The perifusion experiments (dynamic GSIS) were performed using a PERI5 machine (Biorep Technologies, Miami, FL, USA) that allows parallel perifusion of up to 12 independent channels with fully software-controllable inputs via a microfluidic manifold capable of generating customizable media gradients. For each experiment, an estimated 100 IEQ of human islets were handpicked and loaded in Perspex microcolumns between two layers of acrylamide-based microbead slurry (Bio-Gel P-4, Bio-Rad Laboratories, Hercules, CA, USA) by experienced operators. Perifusing buffer containing 125 mM NaCl, 5.9 mM KCl, 1.28 mM CaCl_2_, 1.2 mM MgCl_2_, 25 mM HEPES, and 0.1% bovine serum albumin at 37°C with selected glucose or KCl (25 mM) concentrations was circulated through the columns at a rate of 100 μL/min. After 60 minutes of washing with low glucose (3 mM, G3) solution for stabilization, islets were stimulated with the following sequence: 8 min of low glucose, linear increases of 4, 8, 12, 16, or 20 min to high glucose (11 mM, G11) ran in parallel for a total of 25 min, 15 min of low glucose, 10 min of KCl, and 10 min of low glucose. Samples (100 μL) were collected every minute from the outflow tubing of the columns in an automatic fraction collector designed for a multi-well plate format. The islets and the perifusion solutions were kept at 37°C in a built-in temperature-controlled chamber while the perifusate in the collecting plate was kept at <4°C to preserve the integrity of the analytes. Insulin concentrations were determined with commercially available human ELISA kits (Mercodia Inc., Winston Salem, NC, USA). Values obtained with the human kit were converted from mU/L to μg/L using 1 μg/L = 23 mU/L per the manufacturer guidelines. Because accurately assessing islet mass in islet equivalent (IEQ) units is nontrivial [36, 37], to account for possible differences among islets in different channels, values were adjusted based on the response to KCl using the area under the curve (AUC) in each column for normalization as described before [18, 35, 38, 39]. All responses are scaled to 100 IEQ.

For calculation purposes, we considered the instantaneous “square wave” increase to occur over a 2 min average as it arrives to the β-cells distributed throughout the islets – some of this due to delays and mixing of the solution as it travels along the tubing of the apparatus and some due to diffusional delays within the isolated and thus unvascularized islets. The latter is more likely to cause noticeable delay in the outflow of insulin than in the inflow of glucose as the diffusion time to the core of an islet, which is calculated as ratio between the square of the distance and the diffusion coefficient, *L*^2^/*D* [40], is around 10 s for glucose and 100 s for insulin assuming a standard islet with a diameter of 150 μm. Plus, for such spherical structures, most of the cells are located closer to the surface than to the core making the average delay even less than these estimates.

### Statistical analysis

Data used here are averages of four samples for each condition. All graphic preparations and curve fittings if needed were performed using GraphPad Prism (GraphPad, La Jolla, CA, USA).

### Computational model

The computational model used here is a slightly updated version of our local concentration-based insulin secretion model implemented within a finite element method (FEM) framework in COMSOL Multiphysics (Figure 5) [34, 35]. Briefly, four concentrations are fully modeled for convective and diffusive mass transport including consumption or release (i.e., reaction) rates: glucose, oxygen, insulin, and “local” insulin (*c*_g_, *c*_o_, *c*_i_, and *c*_iL_). Inclusion of a “local” insulin compartment in the model was needed for a correct time-scale of insulin release, so that insulin is assumed to be first secreted into this compartment and then released from there following first order kinetics, 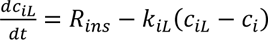 (Figure 5) [34].

#### Mass transport and consumption rates

Diffusion is described by eq. 1 being the generic diffusion equation (nonconservative formulation, incompressible fluid)

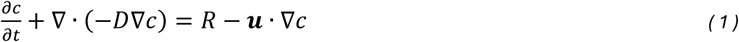

where, for each species, *c* denotes concentration [mol·m^−3^], *D* diffusion coefficient [m^2^·s^−1^], *R* reaction rate [mol·m^−3^·s^−1^], ***u*** the velocity field [m·s^−1^], and ∇ the standard *del* (*nabla*) operator 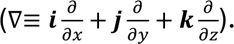 All diffusion coefficients (*D*) used here are from the previous model [34, 35]. All consumption and release rates are assumed to follow Hill–type sigmoid dependence on the corresponding local concentrations:

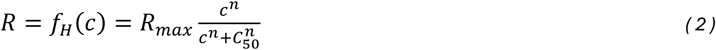

In all cases (i.e., insulin, glucose, oxygen), *R*_max_ denotes the maximum reaction rate [mol·m^-3^·s^-1^], *C*_50_, the concentration corresponding to half-maximal response [mol·m^-3^], and *n*, the Hill slope characterizing the shape of the response. Parameter values (*R*_max_, *C*_50_, etc.) are the same as in the original model, except for those re-calibrated and explicitly discussed here. For oxygen and glucose consumption, the parameter values used in our previous model [34, 35] were maintained (*R*_max,oxy_ = –0.034 mol·m^-3^·s^-1^, *C*_50,oxy_ = 1 μM; *R*_max,gluc_ = –0.028 mol·m^-3^·s^-1^, *C*_50,gluc_ = 10 μM) including Hill coefficients of unity (*n*_oxy_ = *n*_gluc_ = 1) as the assumption of regular Michaelis-Menten kinetics gave adequate fit. Oxygen consumption is assumed to consist of a base-rate and an additional component that increases due to the increasing metabolic demand in parallel with the insulin secretion rate. Sections where the oxygen concentration fells below a critical value of *C*_cr,oxy_ = 0.1 μM are assumed to become necrotic and no longer functional; for additional details see [34].

#### Insulin secretion rates

A main part of the model, and the only one recalibrated here, is the functional form describing the glucose-dependence of the insulin secretion rate, *R*_ins_. *R*_ins_ is assumed to be the sum of a second-phase release, related to the local glucose concentration (*c*_g_) that reaches the β-cell, and a first-phase release, related to its rate of change (∂*c*_g_/∂*t*), with both being described by Hill-type functions as shown below in eq. 3 and 4. Even if glucose is not directly an enzyme substrate for insulin production and thus there is no direct justification for the use of Michaelis-Menten–type kinetics, the Hill equation, a generalized form of the Michaelis-Menten one that has *n* = 1, was found to provide a functionality that fits the experimental results with high accuracy. Accordingly, the main function used here to describe the glucose-insulin dynamics of the second-phase response is eq. 3 as shown below:

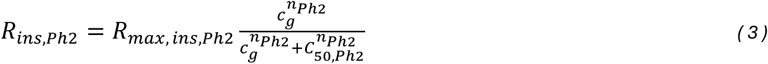

with *n*_Ph2_ = 3.2, *C*_50,Ph2_ = 7.9 mM, and *R*_max,ins,Ph2_ = 1.8×10^-5^ mol·m^-3^·s^-1^ (slightly rescaled from the original model, which had *n*_Ph2_ = 2.5, *C*_50,Ph2_ = 7.0 mM, and *R*_max,ins,Ph2_ = 3.0×10^-5^ mol·m^-3^·s^-1^, to match our more recent experimental concentration dependency of second-phase responses [18]). Note, however, that with the Hill function as used here (eq. 3), insulin secretion tends toward 0 as the glucose concentration decreases, whereas stressed islets are known to leak insulin even at basal glucose levels [39].

The first-phase response is incorporated via a component that depends on the rate of glucose increase, i.e., the glucose time-gradient (*c_t_* = ∂*c*_gluc_/∂*t*). This is non-zero only when the glucose concentration is increasing, i.e., only when *c*_t_ > 0. Again, a Hill–type sigmoid response was assumed to ensure a plateau:

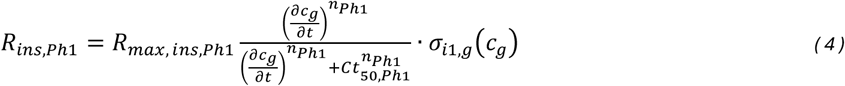

with *n*_Ph1_ = 2.0, *Ct*_50,Ph1_ = 0.0225 mM·s^-1^ (1.35 mM/min), and *R*_max,ins,Ph1_ = 9.0×10^-5^ mol·m^-3^·s^-1^ – values that were somewhat rescaled now compared to the original model to better fit the first-phase experimental data obtained here. With the *Ct*_50_ value used in the model here (1.35 mM/min), the sigmoid response function gives an approximately linear response for a rate range that covers most of the normal physiologic conditions (e.g., 3 to 6 mM increase in 15–30 min: 0.1–0.4 mM/min) [16] and well beyond (up to about 1.5 mM/min; Figure 4B). A fully linear (i.e., proportional) glucose gradient dependent term has been used in several models including for artificial pancreases mainly following Jaffrin [41-45].

*R*_ins,Ph1_, as used here (eq. 4), also incorporates one additional modulating function *σ*_i1,g_ to reduce this gradient-dependent response for islets that are already operating at an elevated second-phase secretion rate and to maximize it around *c*_g_ values where islets are likely to be most sensitive (*C*_m_ = 5 mM). The corresponding function used here (eq. 5) was slightly modified from its previous form [34] and is now the product of a bell-shaped curve (derivative of a sigmoid function) and a sigmoid cut-off function (*C*_cut_ = 15 mM):

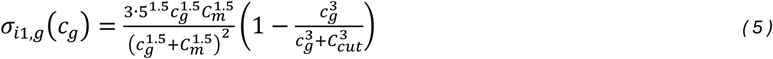

Finally, total insulin release is obtained as the sum of the first- and second-phase releases (plus an additional modulating function to account for the limiting effect of oxygen availability; see [34] for details):

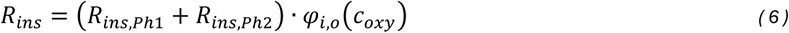

#### Fluid dynamics model

To account for media flow in the perifusion chamber, the convection and diffusion models need to be coupled to a fluid dynamics model, and this is done via the incompressible Navier–Stokes model for Newtonian flow (constant viscosity):

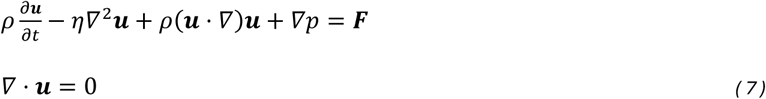

Here, ***u*** is the velocity field, *ρ* denotes density [kg·m^-3^], *η* viscosity [kg·m^-1^·s^-1^ = Pa·s], *p* pressure [Pa, N·m^-2^, kg·m^-1^ ·s^-2^], and ***F*** volume force [N·m^-3^, kg·m^-2^·s^-2^]. The first equation in eq. 7 is the momentum balance; the second one is simply the equation of continuity for incompressible fluids.

#### Model implementation

The model was implemented in COMSOL Multiphysics 6.1 (COMSOL Inc., Burlington, MA). The time-dependent (transient) solver was used with intermediate time-steps. Computations were performed with the MUMPS direct solver as linear system solver and with an enforced maximum step of 0.5 s to ensure that changes in the incoming glucose concentrations are not missed due to overstepping of the solver. For the parameters rescaled here, parametric sweeps were used to cover all the different glucose ramps and steps in one run, and optimal parameter values obtained following iterative narrowing of the swept range were then selected based on minimizing the sum of squared errors (SSEs) between predicted and measured insulin secretion rates.

As before, a representative case, a 2D cross-section of a cylindrical tube with two spherical islets of 100 and 150 μm diameter was used (see Figure 7 for illustration). The constant ramps for increase in the incoming glucose concentration were implemented using the smoothed Heaviside step function at predefined time points *t_i_*, with adjustable transition time *τ*: *c*g = *c*low + *c*step·flc1hs(*t* – *ti*, *τ*). To account for the unavoidable mixing in the tubing and diffusional delays in reaching the β-cells of the islets, the glucose ramps were considered 1 min longer for modeling than the actual setup in the perifusion machine, i.e., 9 min for the 8 min ramp. For FEM, COMSOL’s physics-controlled ‘Finer’ mesh size was used (∼10,000 mesh elements). Boundary, inflow, and outflow conditions were the same as defined in the original model [34, 35]. The insulin outflow flux from the COMSOL 2D model, which is in mol/m/s units, was converted to the units used in the figures (pg/min/IEQ; equivalent to the μg/L concentration measured with the 100 IEQ per column and the 100 μL/min flow rate used for perifusion) by multiplying with the height needed for the corresponding IEQ volume and the molecular weight of insulin then dividing with the outflow volume per minute (2.41×10^13^). Since scaling to IEQ is somewhat arbitrary, the COMSOL-calculated values shown to illustrate the agreement with the experimental values (Figure 6) were scaled by 0.78 to optimize overlap.

## Results

We performed dynamic perifusion studies with isolated human pancreatic islets using a system that allowed adjustable transition times for the increase in incoming glucose concentrations and established the corresponding insulin secretion time-profiles with once a minute time-resolution. These confirmed the presence of a first-phase peak that decreased as the gradient (the rate of increase) of the incoming glucose concentration decreased (Figure 2). Studies were performed using a fully automated machine (PERI5) with software-controlled customizable inflow for up to 12 independent channels and outflow collection with adjustable temporal resolution (1 min used here). This way, islets from the same batch could be perifused in parallel to allow a direct comparison of the differences in the responses due to the different rates of change in the incoming glucose. Studies were performed with a glucose step of 8 mM (from 3 to 11 mM), the standard used in most of our previous studies here at the Diabetes Research Institute (DRI). This allowed us to include considerable historical data for the corresponding “square wave” glucose step as an average for reference (Figure 1) [18, 35, 46, 47]. Here, however, in addition to the abrupt “square wave” increase, we also used gradual increases over times of 4, 8, 12, 16, and 20 min in parallel channels resulting in glucose gradients in the 0.4–4.0 mM/min range (Figure 2).

**Figure 2.**
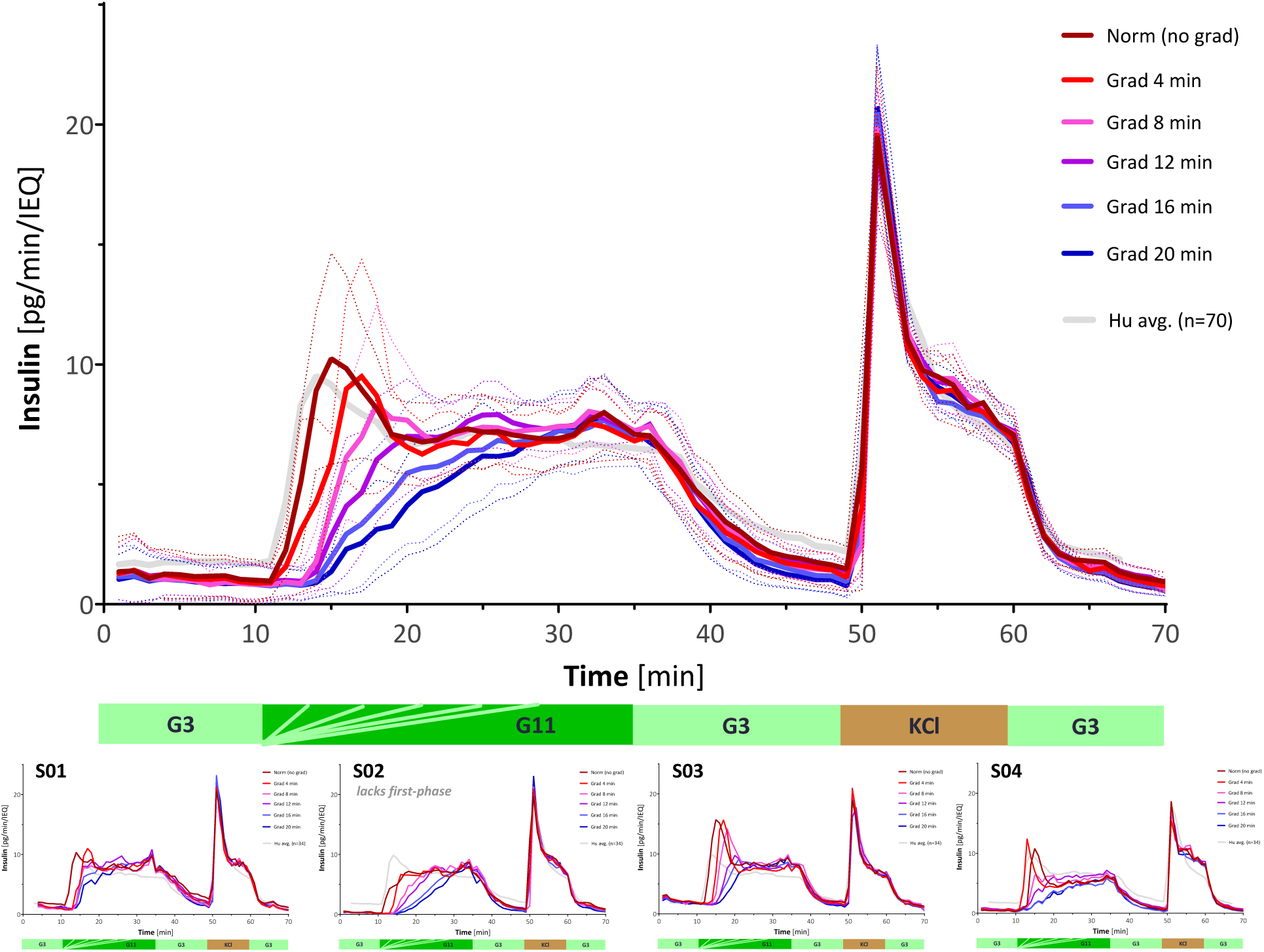
The effect of the rate of glucose increase on the insulin secretion of human islets. Summary of experimental data for isolated human islets perifused in parallel using different rates of glucose increase (3 mM increased to 11 mM in 0, 4, 8, 12, 16, and 20 min as indicated, followed by 15 min low 3 mM glucose, 10 min 25 mM KCl, and 10 min low 3 mM glucose; corrected to ∼100 IEQ per chamber; *n* = 4 per group). Data (insulin secreted per minute) are shown color-coded from red (4 min) to blue (20 min); the average of all data obtained in our lab for the stepwise G3→G11 protocol (no gradient, *n* = 70; same as shown in Figure 1) is included as a dashed grey line for reference. The individual responses obtained for each of the four different islet isolations (all from normal non-diabetic subjects; S01–S04) are included at the bottom to illustrate the variability; one (S02) clearly lacked first-phase response.

For a quantitative assessment of the first-phase insulin release, we defined it to be the excess amount of insulin secreted in addition to what would be expected from the glucose concentration-dependent second-phase release (Figure 3A) and plotted it as a function of the time (Figure 3B). It is generally accepted that β-cells sense and respond both to the glucose concentration itself (*c*_g_) and to its rate of change (gradient; ∂*c*_g_/∂*t*). Here, in line with our computational model [34, 35], we assumed that the rate of insulin release (*R*_ins_) depends on these via two essentially independent processes with a second-phase release related to the local glucose concentration (*c*_g_) that reaches the β-cell and a first-phase release related to its rate of change (∂*c*_g_/∂*t*). Previously, we found the second-phase secretion of perifused isolated human islets to be described very well by a Hill-type sigmoid functions (eq. 3) with a half-maximal concentration of *C*_50,Ph2_ = 7.9±0.3 mM and a Hill slope of *n*_Ph2_ = 3.2±0.4 (*r*^2^ = 0.98) (Figure 4A) [18]. Since this corresponds to an essentially linear response across the blood glucose range of 3 to 11 mM used here, the insulin released during the peak of the first-phase response can be considered as consisting of an essentially concentration-proportional second-phase part that rises in parallel with the increase in the glucose concentration plus an “excess” part due to the rate-responsive first-phase response (Figure 3A). This can be quantified for the different glucose ramps used here resulting in the time-profiles shown in Figure 3B. Plotting the corresponding average secretion rates as a function of the rate of glucose increase (mM/min) results in the rate-dependency shown in Figure 4B. Considering once again that ultimately this response also has to be limited (i.e., plateauing out at a maximum), it was also fitted with a sigmoid function similar to the one used for the second-phase response just with ∂*c*_g_/∂*t* and not *c*_g_ as its argument (eq. 4) resulting in a reasonable fit (*r*^2^ = 0.96) with a half-maximal rate of *Ct*_50,Ph1_ = 1.24±0.1 mM/min and a Hill slope of *n*_Ph1_ = 1.8±0.3 (Figure 4B). The idea that the first-phase response is the excess over the essentially concentration-proportional second-phase part is also supported by the lucky coincidence that islets from one of the four subjects assayed here seem to lack this response (Figure 2, S02) despite all islets being from normal non-diabetic donors.

**Figure 3.**
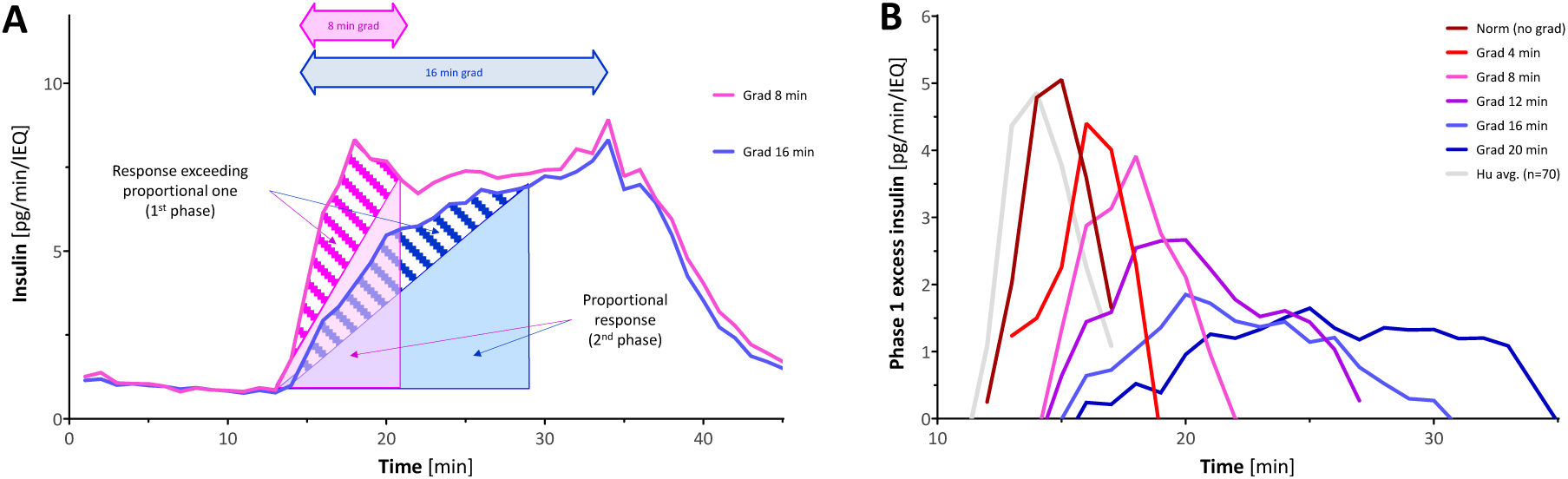
Amount of first phase “excess” insulin released beyond that expected based on a concentration-dependent (second-phase) response only. (**A**) Procedure used to calculate the amount of insulin in excess of the concentration-dependent proportional one (shaded area on top of the colored triangles; illustrated here for the 8 and 16 min G3 to G11 increases from Figure 2). (**B**) Time profile of the insulin that can be considered as the rate-responsive (phase 1) part released in excess of the concentration-dependent (phase 2) part as calculated with the procedure shown in A. Color coding same as in Figure 2.

**Figure 4.**
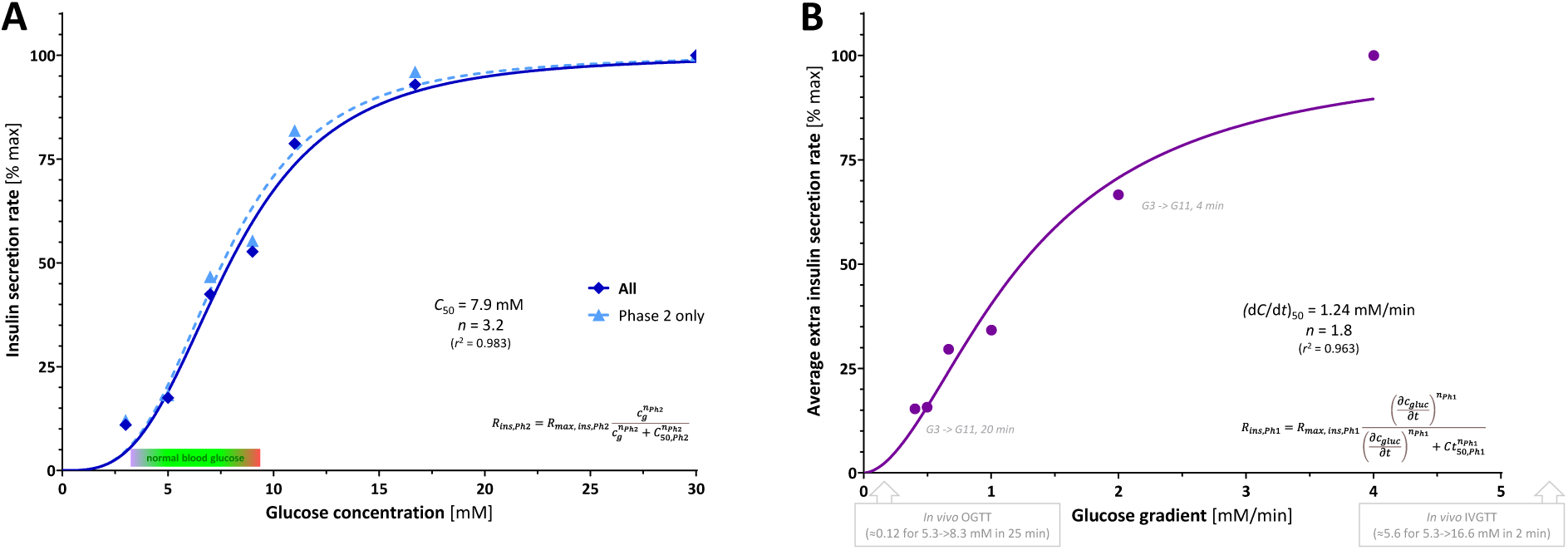
Dependence of insulin secretion on glucose concentration and its increase rate. (**A**) Average total (all) and phase 2 insulin secretion rates at different glucose concentrations (symbols; data expressed as percent of maximum at G30; values from [18]) fitted with Hill-type sigmoid functions (line). Best fit for phase 2: slope *n*_Ph2_ = 3.2±0.4 and half-maximal concentration *C*_50,Ph2_ = 7.9±0.3 mM (*r*^2^ = 0.98). (**B**) Average phase 1 insulin secretion for different glucose increase rates as calculated in Figure 3 (symbols; expressed as percent of maximum) and fitted with a Hill-type sigmoid function (line). Best fit for phase 1: slope *n*_Ph1_ = 1.8±0.3 and half-maximal concentration gradient *Ct*_50,Ph1_ = 1.24±0.12 mM/min (*r*^2^ = 0.96).

Next, as a further proof that this mechanism is realistic, we used these first-phase experimental responses to fine-tune the calibration of the corresponding part of our previously developed computational model of glucose-insulin control system relying on local concentrations and built on a general framework of a sigmoid proportional–integral–derivative (σPID) controller (Figure 5) [34, 35]. The model needed a slight recalibration of its first-phase parametrization to improve fit to the present data (see Methods for details). The resulting recalibrated model was able to give good description of both the first- and second-phase responses in our perifusion assays designed to evaluate the effects of both increasing glucose concentrations (Figure 6B; experimental data from [18]) and increasing rates of increase (Figure 6A; present data). Agreement between the model predicted and the experimental data measured here is quite good – both the shapes and the trends of the time-profile curves are well predicted by the model (Figure 6A) accounting for 90% of the variability in the data (*r*^2^ = 0.90; a plot of model predicted versus observed experimental data is shown in Supporting Figure S1A). It is also good for the time profiles measured with different glucose steps earlier [18] (Figure 6B and Figure S1B; *r*^2^ = 0.91), which were included as they also represent different rates of increase; only the first-phase bump of the largest step is not predicted well. Finally, a 2D graph is also included for illustration depicting the calculated insulin production during first phase versus that in second phase in the two sample islets used for modeling (Figure 7A,B) and a corresponding 3D illustration with the insulin as height data (Figure 7C,D).

**Figure 5.**
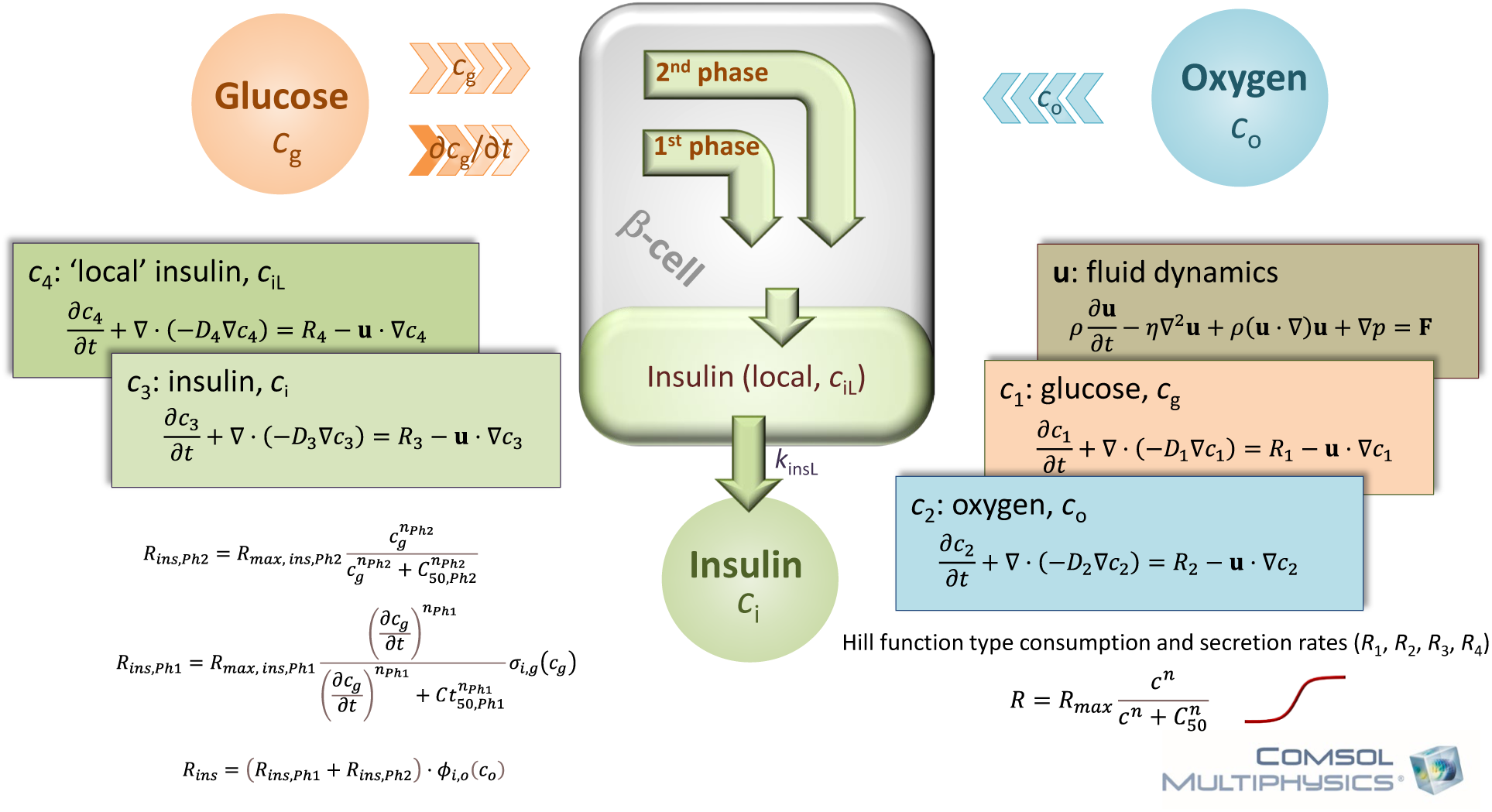
Schematics of the computational model of insulin secretion utilized in this study. Insulin secretion is expressed as the sum of second- and first-phase secretions determined by the local glucose concentration, *c*_g_, and its rate of change, ∂*c*_g_/∂*t*, respectively via Hill-type sigmoid functions (plus also modulated by the availability of local oxygen, *c*_o_).

**Figure 6.**
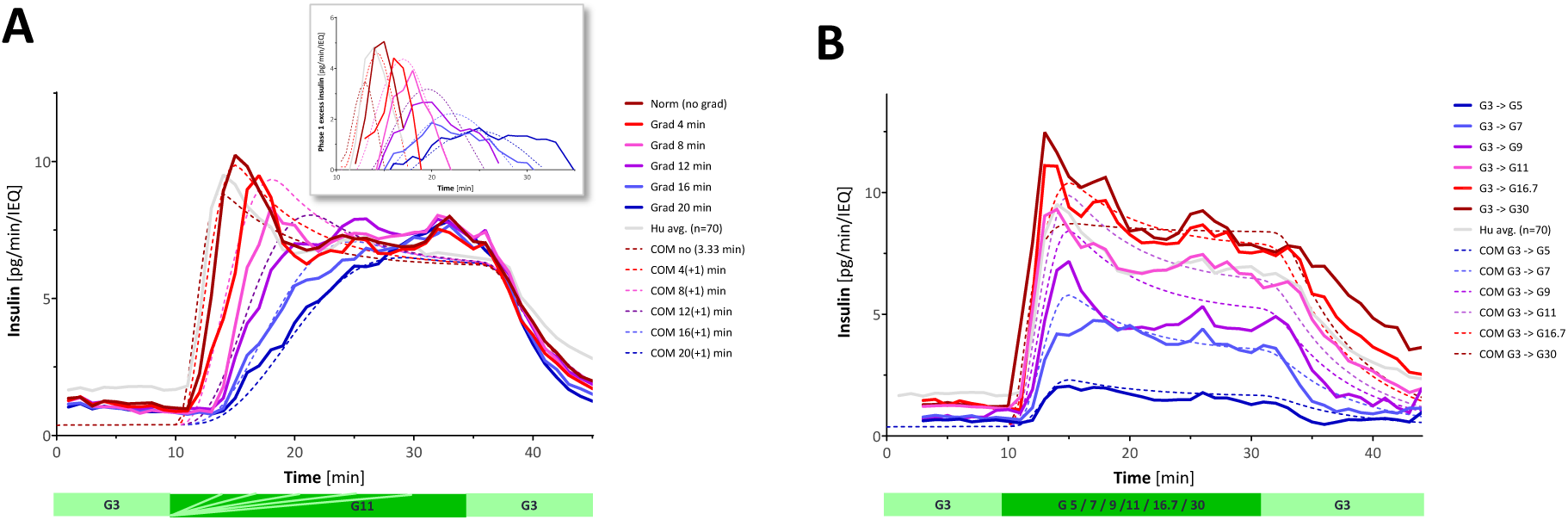
Dynamic perifusion GSIS time profiles and corresponding predicted values with the present COMSOL Multiphysics computational model. Experimental time profiles (continuous lines color-coded from dark red to blue) obtained in the present different rates of increase experiment (Figure 2) (**A**) and in our previous different glucose steps experiment [18] (**B**) superimposed with corresponding predictions of the COMSOL model as recalibrated here and scaled by 0.78 to optimize overlap (dashed lines with the same color code). Inset in A shows the same for the first-phase excess insulin only. Corresponding plots of predicted versus observed data are shown in Supporting Figure S1.

**Figure 7.**
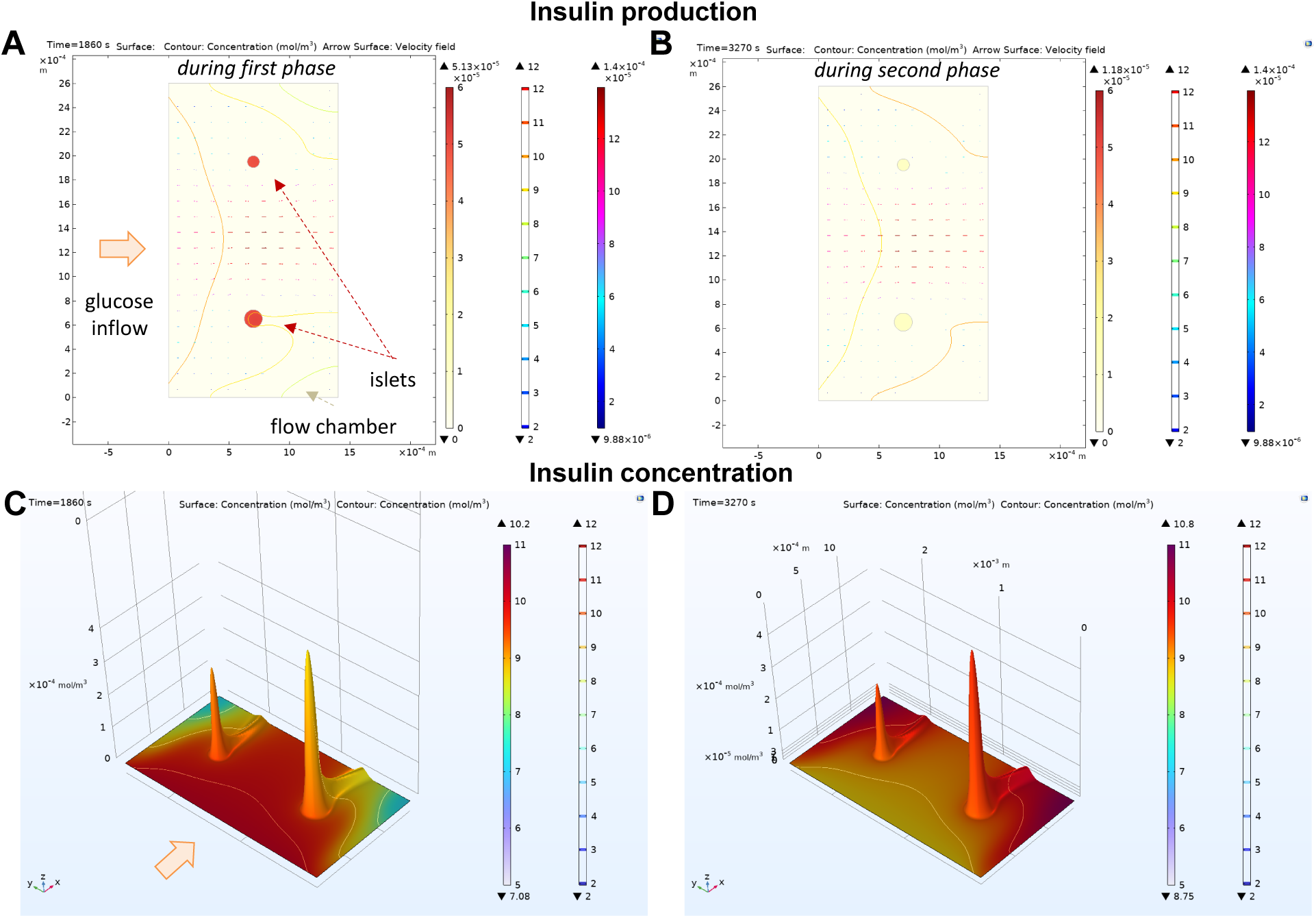
Model-calculated insulin concentration and production rates during first- and second-phase responses. (**A**, **B**) Calculated insulin production rates for two representative islets of *d* = 100 and 150 μm diameters shown as surface plots color-coded from yellow for low to dark red for high during first phase (A; increasing glucose concentration) and end of second phase (B; start of decreasing glucose concentration) insulin secretion. Figure also shows the flow of the perifusion fluid as streamlines color-coded for velocity and incoming glucose concentrations as iso-levels color coded from blue for low to red for high. (**C**, **D**) Model-calculated insulin concentrations for the same conditions shown as height data with surface plots color-coded for glucose concentration (from blue to red). All data shown are for the perifusion concentrations used here with glucose increasing from 3 to 11 mM under normoxic conditions (atmospheric oxygen; ∼20% ≈ 0.2 mM).

## Discussion

To the best of our knowledge, the present work is the first time that the biphasic nature of GSIS of isolated human islets was assessed in detail *in vitro* using various gradual changes in the incoming glucose concentration. To attempt to quantify the rate-dependence of the first-phase response, limited numbers of constant glucose ramps have been explored already by Grodsky in 1972 using perfused rat pancreas (225 mg/dL increase in 5 and 60 minutes corresponding to 2.5 and 0.21 mM/min, respectively) and by Nomura in 1984 with perifused rat islets (0.09–0.26 mM/min) [41]. However, these were too small gradients to allow a clear separation between first- and second-phase responses for quantitation and were further confounded by the rising second-phase responses of rat islets. With the 8 mM step used here, glucose increase rates evaluated by us ranged from 0.4 to 4.0 mM/min. This covers a reasonable range for perifusion experiments for the present setup as well as the range between the clinically used oral and intravenous glucose tolerance tests (OGTT and IVGTT, respectively; Figure 4B), which were found to be ∼0.12 mM/min in OGTT (blood glucose increasing from 5.3 mM to 8.3 mM in ∼25 min) and ∼5.6 mM/min in IVGTT (blood glucose increasing from 5.3 mM to 16.6 mM in 2 min) [16].

Admittedly, square wave stimulations such as those typically used to elicit first-phase responses are not physiological as such abrupt changes in blood glucose do not occur normally since blood glucose levels rise more gradually following oral food intake *in vivo*. However, while a clear first-phase insulin release may not be obvious after an oral meal or glucose challenge as the corresponding increase in glucose is not sufficiently abrupt, it has been shown to occur *in vivo* following intravenous glucose tolerance tests (IVGTTs) [16, 48] or in hyperglycemic clamp studies [49-51]. Notably, the biphasic pattern is considered a real characteristic [13] and a sensitive indication of adequate β-cell function [19, 20]. There is evidence that the ability of β-cells to generate a rapidly increasing insulin profile (first-phase response) is important for achieving optimal interstitial insulin concentrations [52] and for restraining hepatic glucose production [19]. First-phase insulin response has been found to be preferentially damaged in the early stages of type 2 diabetes (T2D) [13] (and references therein). Its accelerated loss in those progressing toward T1D has also been shown in several studies [22-26]. Because insulin is secreted directly into the portal system, there is greater hepatic than peripheral insulinization making the liver the most likely target of first-phase insulin secretion, and even under physiological conditions where a biphasic pattern is not produced, the early insulin response may be aimed at rapidly shifting glucose metabolism from fasting to prandial state by primarily acting on the liver [21]. In fact, it has been hypothesized that the dynamic properties of the β-cell that generate the first-phase response and hence the biphasic pattern in response to a rapid glucose infusion (e.g., IVGTT) could be the same as those operating with gradual entry of glucose from the gut (e.g., OGTT) [21]. It has also been suggested that some measures derived from OGTT correlate with first-phase insulin response and have predictive ability for T1D [53]. Hence, inclusion of a physiologically accurate first-phase insulin response is important for artificial or bioartificial pancreas devices since its absence could lead to long-term physiological consequences [35, 54].

We found the GSIS of perifused human islets to be well-describable as the sum of a rate-sensitive first phase, which is a sigmoid function of the glucose gradient with half-maximal value around 1.25 mM/min, and a concentration-sensitive second phase, which is a sigmoid function of the glucose concentration with half-maximal value around 8 mM (Figure 4). Since the second-phase secretion of perifused isolated human islets seems to be flat even for up to one hour [18] (versus a rising one in rodents, especially rat islets – Figure 1 and [55]), a straightforward concentration-dependence can be used to describe it (i.e., there is no need for a time-dependent part). Because it has to be limited as all biological responses (plateauing out at a maximum – saturation is clearly evident above 16 mM for human insulin secretion [18]), use of a Hill-type sigmoid function such as eq. 3 makes sense [34]. Indeed, we found it to account for more than 98% of the variability in our data (*r*^2^ = 0.98) with *C*_50,Ph2_ = 7.9±0.3 mM and *n*_Ph2_ = 3.2±0.4 (Figure 4A) [18]. Note that this corresponds to an essentially linear response across the blood glucose range of 3 to 11 mM, which was used here as the glucose step (Figure 4A) allowing us the quantification of the first-phase response as shown in Figure 3A. It is also notable that these parameters (*C*_50_ = 7.9 mM, *n* = 3.2) are very similar to those characterizing the function of glucokinase, an enzyme that plays a central role in glucose sensing of β-cells and is also well-described by a Hill-type function (average of published experimental values: *C*_50_ = 7.7 mM, *n* = 1.7) [56, 57]. As mentioned earlier [18], our second-phase estimates (*C*_50_ = 7.9 mM, *n* = 3.2) agree quite well with those from the detailed perifusion study by Henquin and co-workers (*C*_50_ = 6.7 mM, *n* = 2.3) [58]. However, *in vivo* hyperglycemic clamp studies found higher *C*_50_s for the second phase, e.g., 14.3±1.3 mM in [49] and 10.8±1.6 mM in [50] (the latter obtained by fitting the average of data from normal subjects with Hill-type functions as used here, eq. 3). Some OGTT minimal models also use higher values, e.g., 17 mM in [59] based on data from [60]. The difference between the *in vitro* and *in vivo C*_50_ values may be due to metabolic limitations with isolated islets that lack vascularization as well as to the absence of systemic modulators (e.g., the incretin effect, which can be responsible for ∼50% of insulin release after glucose ingestion [21]) and other effects, such as those on insulin clearance and distribution, which are present in the *in vivo* but not in the *in vitro* experiments. The hyperglycemic clamp experiments clearly show a rising second-phase response [49-51], especially at higher glucose concentrations, whereas they remained flat in our perifusion studies with human islets at 11 mM [18].

Use of the various glucose ramps in perifusion studies here allowed us to interrogate whether the occurrence of the first-phase insulin secretion is an inherent characteristic of the islets that is present even when these mini-organs are isolated from the numerous influences that are part of their normal physiological environment (e.g., vascularization, innervation, incretins, etc.). Our results show that a first-phase release is only evident if there is a sufficiently steep increase of incoming glucose and that islets can enter the GSIS second phase directly without the apparent need of a previous first phase (Figure 2). Notably, we observed high variability in the first-phase insulin secretion rates among our four subjects, despite all being non-diabetic (Figure 2, bottom row) – differences in donor age, sex, BMI, HbA1c as well as ischemia and culture time and isolation center (Supplementary Table S1) might all contribute. One subject (S02) showed an essentially lacking first-phase response; this subject had the most elevated HbA1c value (5.8%) and longest culture time (86 h), which could be potential causes, but with such a small sample size, this has only observational value. To assess what percentage of healthy subjects might lack a significant first-phase response, we reanalyzed our existing human perifusion data (*n* = 70 control samples) and found that the first-phase peak is highly variable and about 10% of the subjects assayed using our stepwise glucose challenge had a first-phase maximum that was less than 20% higher than the second-phase plateau (Supplementary Figure S2). In agreement with these *in vitro* observations, there is evidence for high variability in the *in vivo* first-phase response as well: while IVGTT is highly reproducible within subjects and can be used to obtain first-phase insulin secretion rates, there is more than six-fold variance between subjects [16, 61].

The glucose-insulin control system as assumed here, including in the corresponding computational model (Figure 5), can be considered a sigmoid proportional–integral– derivative controller (σPID) [34], a generalized PID controller. Linear PID controllers are widely used in technical systems to provide control feedback based on the error signal *ε* (the difference between the existing and desired output). As their name suggests, they involve terms that are proportional with the error itself (*ε*; P), its integral (∫*ε*d*t*; I), and its differential (d*ε*/d*t*; D). Thus, the output signal *ψ* of a PID controller at any given time *t* is calculated as:

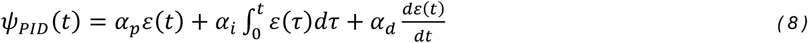

PID controllers have been suggested for the modeling of the glucose-insulin system [1, 62] and especially for closed-loop insulin delivery systems with continuous glucose sensors [3, 63-65]. However, linear (i.e., proportional) responses in biological systems are only possible over a limited range as biological responses, such as enzyme rates, hormone secretion rates, or synapse firing rates, must always plateau at some maximum value. Accordingly, sigmoid, e.g., Hill type functions *f*H (eq. 2) that allow nonlinear responses with limited maximum rates (*R*_max_) and flexible shapes (*n*), are better suited for biological feedback mechanism. Therefore, we suggested sigmoid PIDs (σPID), where terms are proportional to the sigmoid functions (*f*H) of the error signal, its integral, and derivative [34]:

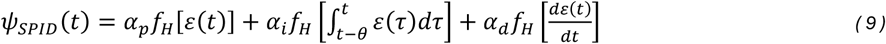

In general, such σPID controllers (eq. 9) should be more suitable to describe biological systems than the widely used linear PIDs (eq. 8) as biological responses are always limited between zero and a possible maximum response. Thus, sigmoid responses are much better suited for their characterization. Note that σPIDs can be considered as essentially linear and providing similar responses as the classic PID control systems, but only across a limited range around their half-maximal points.

The insulin secretion model used here assumes that the second-phase term is proportional with a sigmoid function of *c*_g_ itself (i.e., its deviation from 0, as the “error” signal *ε*; Figure 5) and the first-phase insulin release is related to the sigmoid function of its differential (∂*c*_g_/∂*t* as ∂*ε*/∂*t*). As always, the role of the differential term is to speed up the system, i.e., to give a large correction signal as soon as possible when the monitored value changes suddenly – exactly the role played by the first-phase insulin secretion.

Regarding the limitations of the present model, currently, there is no integral term implemented in it – this will be needed, for example, for responses where there is a rising second phase (e.g., such as those of rat islets; Figure 1). While the actual mechanism of GSIS in β-cells is complex [1, 8-12, 66-69], the present model is just a simplified model intended to account for the main glucose-responsive aspects of insulin secretion; it does not account for several other known important aspects such as the pulsatile / oscillatory nature, time-dependent inhibition and potentiation (e.g., “glucose priming”), or amplifiers such as glucagon-like peptide–1 (GLP-1) [13, 70]. It only focuses on insulin and not on changes in other metabolites along the pathway [71]. It also does not address the specific causes that could result in such sigmoid responses, which could be due to various mechanism involving, for example, reserve and readily releasable pools of secretory granules with differentially distributed response thresholds as has been suggested [1, 11, 13, 72, 73].

## Conclusion

Dynamic perifusion studies with human islets showed that the rate of first-phase insulin secretion increases with the gradient of the glucose increase. If one assumes the first-phase release to be the excess amount released in addition to what would be expected from the glucose concentration-dependent second-phase release, it can be described well by a Hill-type sigmoid function with best fit suggesting a half-maximal increase rate value of 1.25 mM/min and a Hill slope of 1.8 (*r*^2^ = 0.96). These results were used to recalibrate our glucose-insulin computational model built as a sigmoid PID controller assuming essentially independent first- and second-phase responses that are Hill-type (sigmoid) functions of the local glucose gradient and concentration resulting in predictions that are in good agreement with measured first- and second-phase insulin responses.

## Author Contributions

PB originated and designed the project, conceived the study, designed experiments, provided study guidance, analyzed and interpreted data, implemented the computational model, and wrote the manuscript; STC, BW, and OA conducted experiments, collected data, and revised and proofread the manuscript.

## Acknowledgments

The help and support provided by Biorep, Inc. and especially Felipe Echeverry by redesigning and allowing the use of their latest perifusion machines is gratefully acknowledged.

## Supplementary Material

Supplemental material includes Figure S1 showing direct plots of predicted versus observed data for the GSIS time profiles shown in Figure 6; Figure S2 illustrating the variability of insulin secretion as assessed in dynamic perifusion assays with isolated human pancreatic islets; and Table S1, the checklist for reporting human islet preparations used in research.

## Data Availability Statement

The data supporting the conclusions of this manuscript will be made available by the authors upon reasonable requests to any qualified researcher.

## Supplementary Material

### Supplementary Figures

**Supplementary Figure S1.**
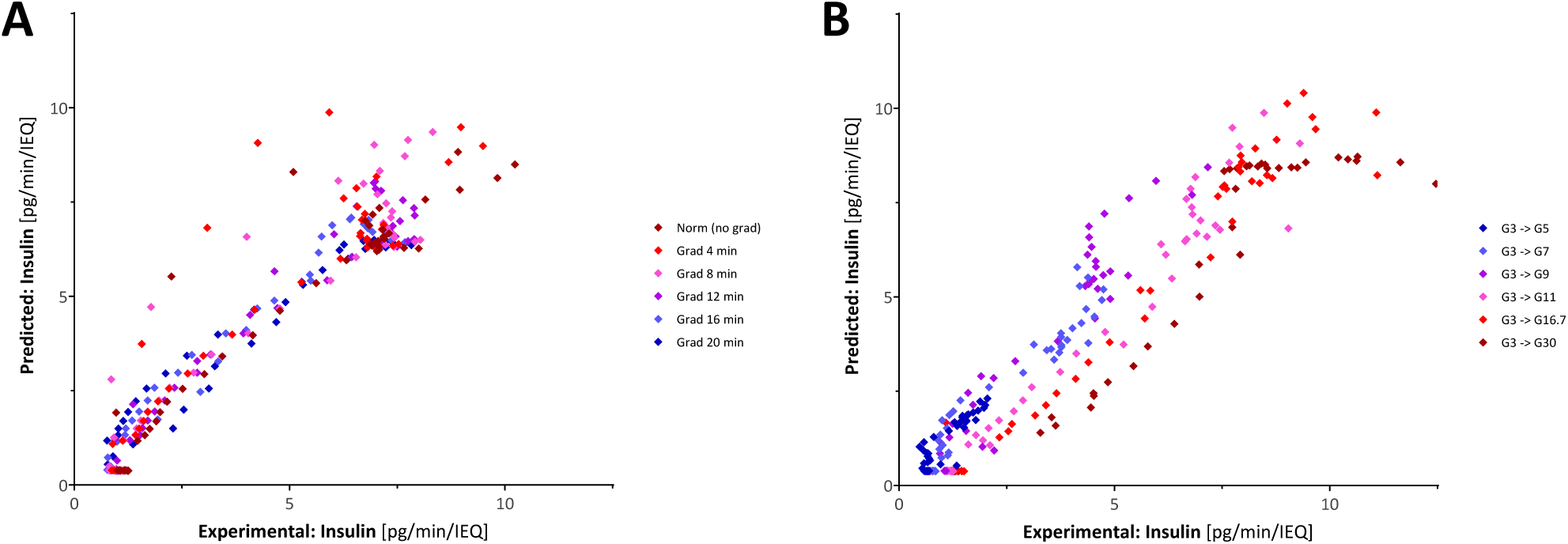
Direct plots of predicted versus observed data for the dynamic perifusion GSIS time profiles shown in Figure 6. Predicted values with the present COMSOL Multiphysics computational model are shown plotted directly versus the corresponding observed experimental data obtained in the different rates of increase (**A**) and different glucose steps experiments (**B**) with the time profiles shown in Figure 6A and 6B, respectively. The model accounts for 90% and 91% of the variability in the data for A and B, respectively (*r*^2^ = 0.90 and 0.91).

**Supplementary Figure S2.**
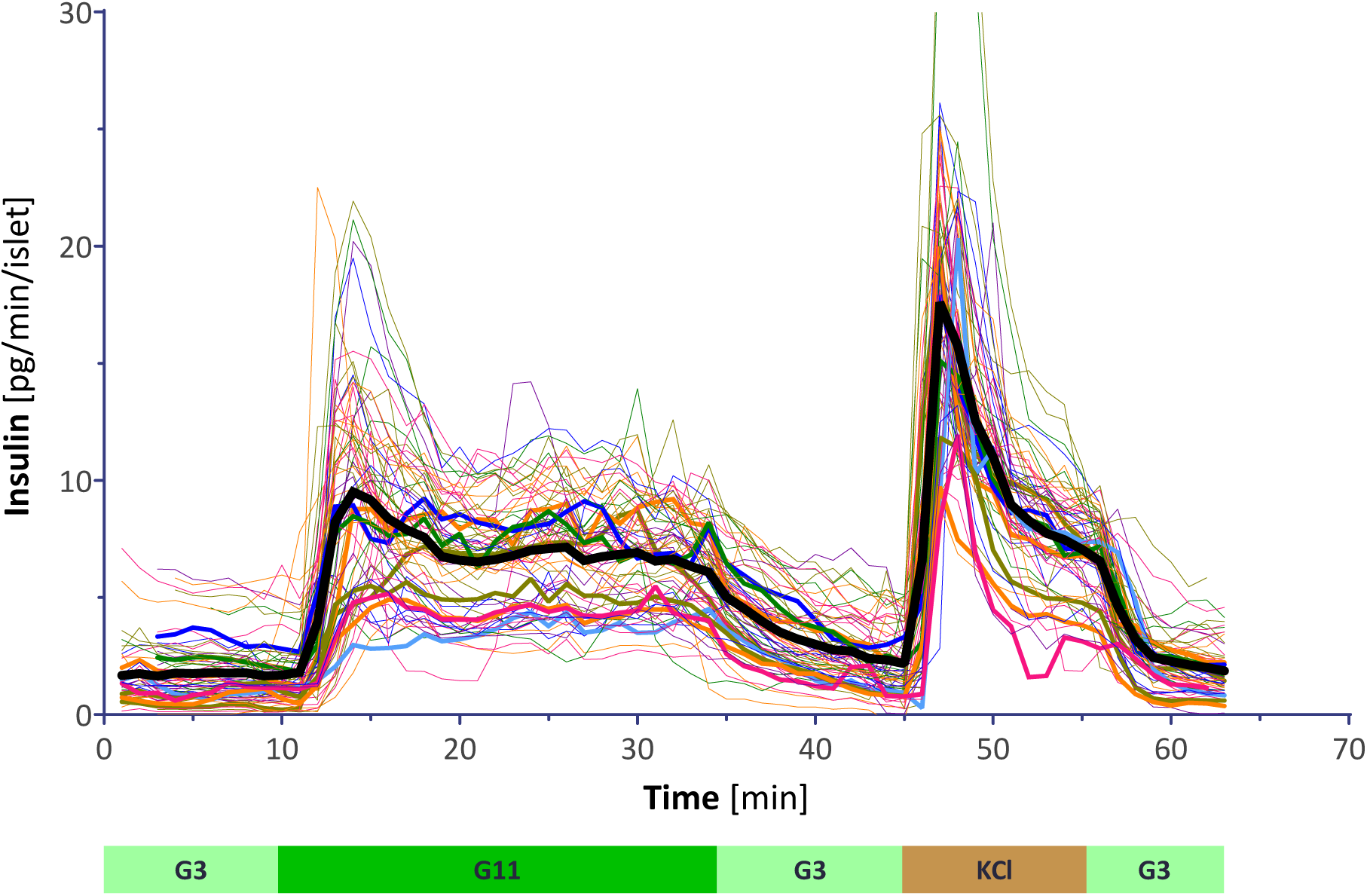
Variability of insulin secretion as assessed in dynamic perifusion assays with isolated human pancreatic islets. Individual time profiles obtained from perifusion assays using our standard protocol (G3→G11→G3→KCl, see Figure 1) shown as spaghetti plot together with their overall average (thick black line; *n* = 70 normal non-diabetic subjects). Data from subjects that showed no significant first-phase response (defined as first-phase maximum <120% of the second-phase plateau) are highlighted with bolder colored lines (*n* = 8).

### Supplementary Tables

**Supplementary Table S1.**
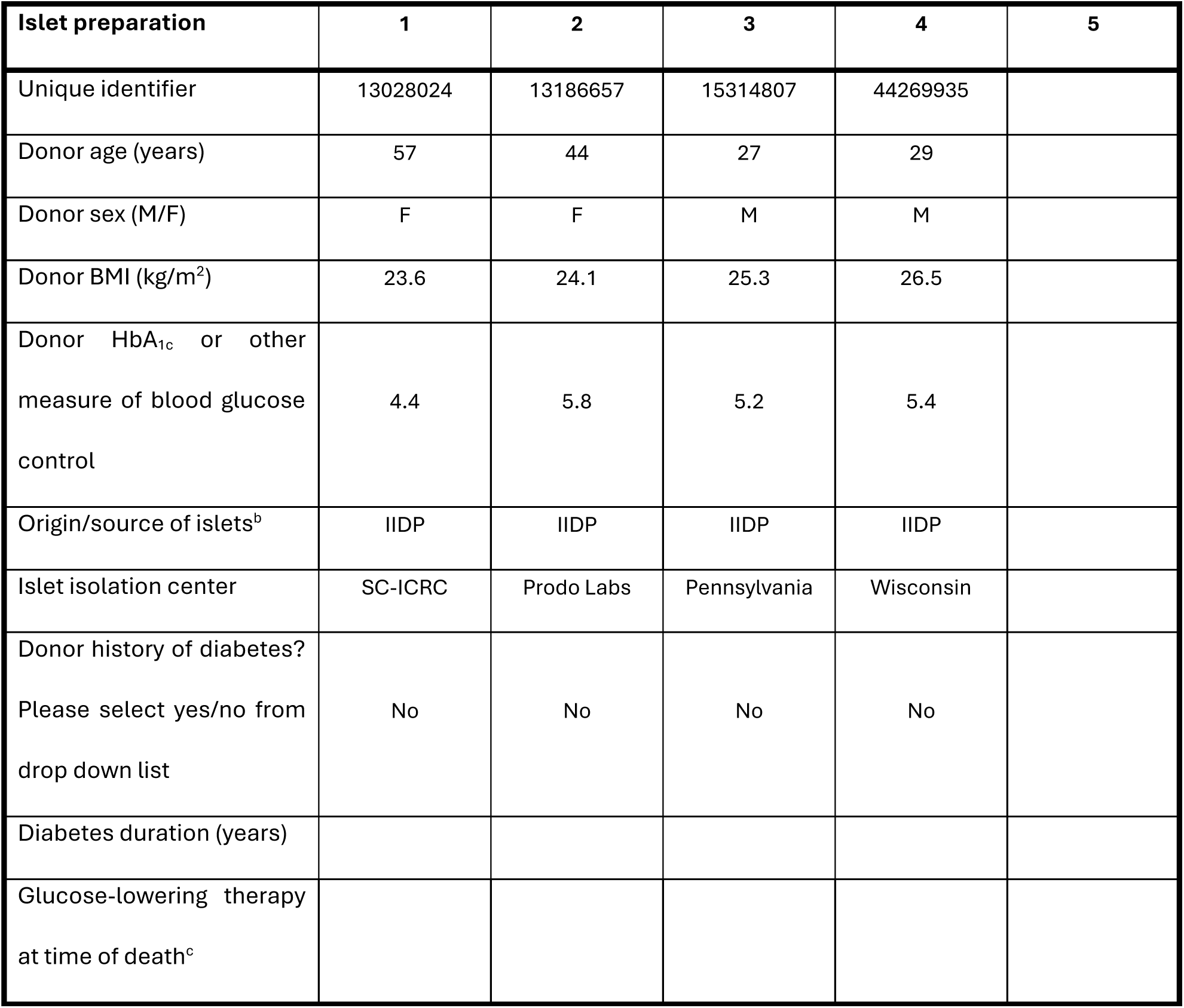
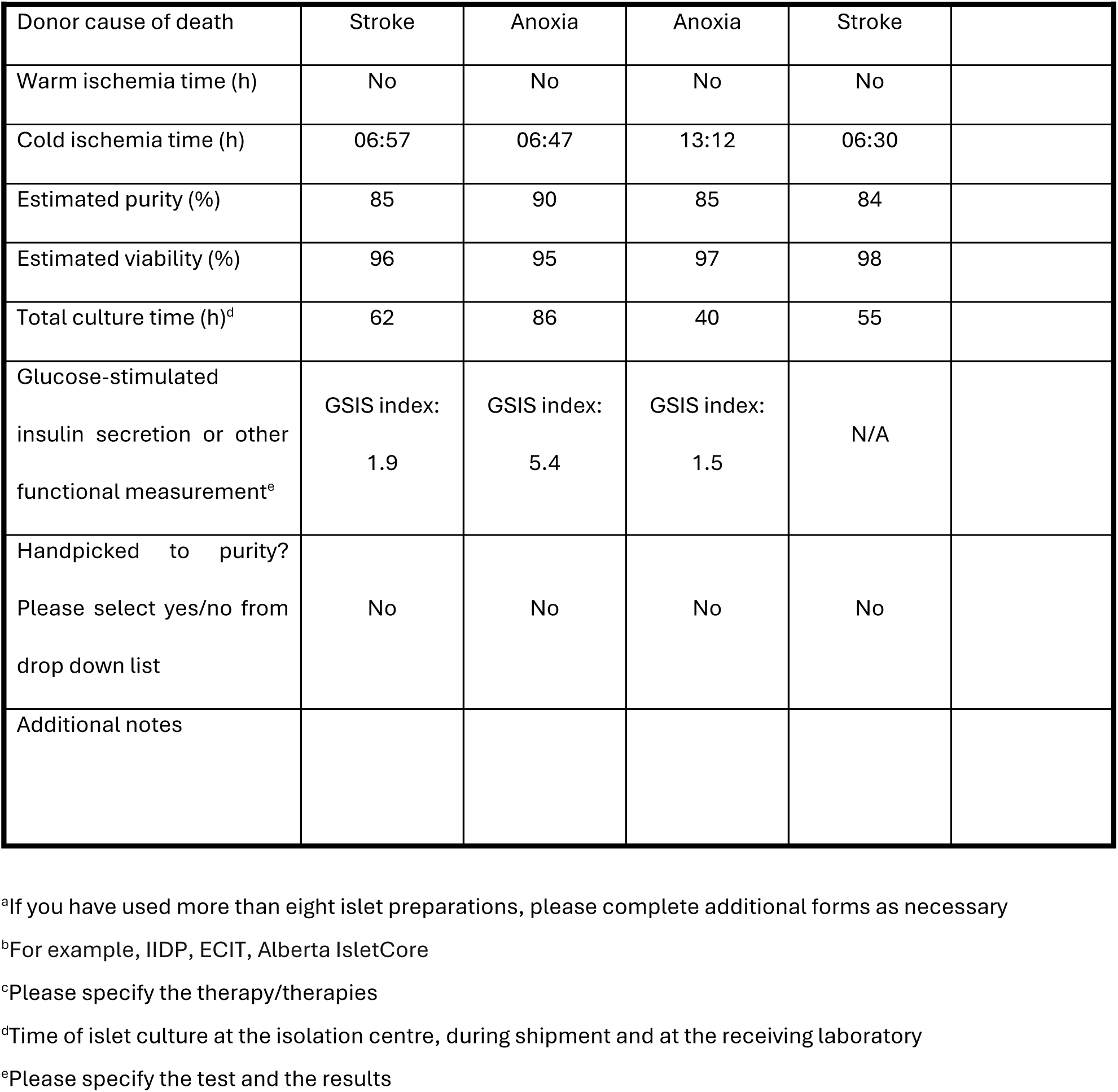
Checklist for Reporting Human Islet Preparations Used in Research. Adapted from Hart NJ, Powers AC (2018) Progress, challenges, and suggestions for using human islets to understand islet biology and human diabetes. *Diabetologia* https://doi.org/10.1007/s00125-018-4772-2.

| Islet preparation | 1 | 2 | 3 | 4 | 5 |
| --- | --- | --- | --- | --- | --- |
| Unique identifier | 13028024 | 13186657 | 15314807 | 44269935 |  |
| Donor age (years) | 57 | 44 | 27 | 29 |  |
| Donor sex (M/F) | F | F | M | M |  |
| Donor BMI (kg/m<sup>2</sup>) | 23.6 | 24.1 | 25.3 | 26.5 |  |
| Donor HbA<sub>1c</sub> or other measure of blood glucose control | 4.4 | 5.8 | 5.2 | 5.4 |  |
| Origin/source of islets<sup>b</sup> | IIDP | IIDP | IIDP | IIDP |  |
| Islet isolation center | SC-ICRC | Prodo Labs | Pennsylvania | Wisconsin |  |
| Donor history of diabetes?<br>Please select yes/no from drop down list | No | No | No | No |  |
| Diabetes duration (years) |  |  |  |  |  |
| Glucose-lowering therapy at time of death<sup>c</sup> |  |  |  |  |  |

|  |  |  |  |  |
| --- | --- | --- | --- | --- |
| Donor cause of death | Stroke | Anoxia | Anoxia | Stroke |
| Warm ischemia time (h) | No | No | No | No |
| Cold ischemia time (h) | 06:57 | 06:47 | 13:12 | 06:30 |
| Estimated purity (%) | 85 | 90 | 85 | 84 |
| Estimated viability (%) | 96 | 95 | 97 | 98 |
| Total culture time (h)<sup>d</sup> | 62 | 86 | 40 | 55 |
| Glucose-stimulated insulin secretion or other functional measurement<sup>e</sup> | GSIS index:<br>1.9 | GSIS index:<br>5.4 | GSIS index:<br>1.5 | N/A |
| Handpicked to purity?<br>Please select yes/no from drop down list | No | No | No | No |
| Additional notes |  |  |  |  |

<sup>a</sup>If you have used more than eight islet preparations, please complete additional forms as necessary
<sup>b</sup>For example, IIDP, ECIT, Alberta IsletCore
<sup>c</sup>Please specify the therapy/therapies
<sup>d</sup>Time of islet culture at the isolation centre, during shipment and at the receiving laboratory
<sup>e</sup>Please specify the test and the results

